# Neural subtypes in developmental stuttering

**DOI:** 10.64898/2026.03.25.714210

**Authors:** Sayan Nanda, Grace Gervino, Cheuk Yee Pang, Emily O. Garnett, Evan Usler, Diane C. Chugani, Soo-Eun Chang, Ho Ming Chow

## Abstract

Developmental stuttering is a complex neurodevelopmental disorder characterized by disfluent speech. At the individual level, the behavioral manifestations of stuttering vary considerably, likely reflecting heterogeneity in underlying neural mechanisms. In this study, we examined individual-specific differences in the brains of children who stutter (CWS), by implementing normative modeling, a framework that quantifies how an individual deviates from an age- and sex-matched reference population. We applied this approach to identify individual-specific structural brain atypicalities using gray and white matter volumes. These volumes were derived from MRI scans from a large mixed-longitudinal dataset of 235 and 240 scans from CWS and fluent controls respectively, aged between 3 and 12 years. Individual deviation maps capturing these atypicalities were then used to cluster CWS into subtypes based on similarities in their neuroanatomical profiles. This analysis identified four neural subtypes with distinct neuroanatomical atypicalities relative to fluent controls. The key findings were a basal ganglia-thalamo-cerebellar subtype associated with higher stuttering severity and lower rates of recovery, and a white matter subtype characterized by mild severity and a higher likelihood of recovery. The remaining two subtypes showed cerebellar differences alongside alterations in brain regions involved in sensorimotor integration. Moreover, cerebellar volume atypicalities were present in all four subtypes, indicating that cerebellar alterations were present across otherwise distinct neural profiles and may represent a shared neuroanatomical feature of stuttering. These findings indicate that examining individual-specific neural differences and subtyping based on patterns of neural atypicalities provides valuable insight into the heterogeneity of developmental stuttering and represents a promising direction for improving our understanding of the disorder.

## INTRODUCTION

Developmental stuttering is a neurodevelopmental disorder associated with frequent, involuntary disruptions in the forward flow of speech, with onset most commonly occurring during early childhood (Smith & Weber, 2017). Speech disfluencies exhibited by people who stutter (PWS) are distinct from typical disfluencies (TD) and are classified as stuttering-like disfluencies (SLDs), consisting of repetitions and dysrhythmic phonations (e.g., blocks, prolongations) (Ambrose & Yairi, 1999). PWS often report a perceived loss of control in their speech, and SLDs are sometimes accompanied by physical concomitants such as face and neck tension, and extremity movements. The disorder affects approximately 5-8% of children and 1% of adults (Månsson, 2000; O. Bloodstein, N. Bernstein Ratner., 2008; Reilly et al., 2009; Yairi & Ambrose, 2005). While the etiology of stuttering is not fully understood, genetic and environmental factors appear to alter the structure and function of neural processes underlying speech motor control in PWS.

At the individual level, the behavioral manifestations of stuttering appear to vary greatly among PWS (Yairi, 2007). This high interindividual variability may reflect a combination of complex etiology underlying stuttering, as well as compensatory or maladaptive responses. To better understand the etiology of stuttering and individualize treatment approaches, efforts have been made to identify subgroups or subtypes of stuttering based on behavioral manifestations, developmental trajectories, temperament, and treatment responsiveness, among others (Ajdacic-Gross et al., 2018; Bloodstein, 1960; O. Bloodstein, N. Bernstein Ratner., 2008; Seery et al., 2007; Yairi, 2007). However, these behavioral subtypes are still not well accepted and have yet to be implemented in a clinical setting. Furthermore, potential subtypes have often been defined based on a limited number of signs or symptoms (Yairi, 2007).

The behavioral heterogeneity observed in stuttering likely reflects differences in underlying neural bases. Consistent with this notion, neuroimaging studies have identified alterations in a wide range of cortical and subcortical regions across studies. While some of this variability may arise from methodological factors, such as small sample sizes and differences in demographics or analytic approach, it may also reflect an inherent neurological heterogeneity within PWS. If stuttering consists of multiple subtypes with distinct neuroanatomical patterns, differences in sampling of these subtypes across studies may contribute to the wide range of neural alterations reported in the literature.

Additionally, some theoretical accounts also propose the existence of neural subtypes of stuttering. A recent review of neuroimaging and neurocomputational modeling work suggested that brain atypicalities in PWS may fall into three broad categories based on the locations of impairments, 1) the basal-ganglia proper, 2) projections between the cerebral cortex, basal ganglia and thalamus, 3) and cortical regions that process cognitive and sensorimotor aspects of speech (Chang & Guenther, 2020). Additionally, empirical evidence for potential neural subtypes of stuttering has been shown based on various neuroimaging measures, such as cerebral lateralization, motor and premotor cortical thickness, and functional connectivity (Chang et al., 2018; Chow et al., 2023; Garnett et al., 2018; Rowe et al., 2024). However, subtyping based on brain function and structural atypicalities in PWS is still in its early stages.

Converging evidence suggests that stuttering is associated with deficits in motor control neural networks (Chang et al., 2019; Neef & Chang, 2024). Recent neuroimaging studies have shown structural and function alterations in the basal-ganglia thalamocortical (BGTC) loop, which supports the precise timing control of the initiation and termination of motor programs (Chow et al., 2023; Frankford et al., 2021; Giraud et al., 2008; Neef et al., 2018; Watkins et al., 2008). In addition to the BGTC loop, atypical structure and function have been reported in sensorimotor cortical regions including the motor, premotor and auditory cortices, inferior frontal gyrus, insula, and the temporoparietal junction, that transform sensory speech representations into coordinated articulatory plans (Fox et al., 1996; Hickok & Poeppel, 2007; Watkins et al., 2008). Consistent with this, a number of diffusion tensor imaging studies of PWS show reduced white matter integrity between left posterior and anterior cortical speech motor areas, including the superior longitudinal fasciculus (SLF), arcuate fasciculus (AF) and frontal aslant tract (FAT) (Cai et al., 2014; Chow & Chang, 2017; Connally et al., 2014; Cykowski et al., 2010; Kell et al., 2009; Watkins et al., 2008). Disruptions in these cortical and white matter pathways may provide unstable or inefficient input to the BGTC loop, affecting speech initiation and sequencing (Hickok et al., 2011; Max et al., 2004).

In addition to the basal ganglia, the cerebellum is a key structure involved in generating detailed motor programs and predicting sensory outcomes through forward models, as well as refining these programs through adaptive adjustments and real-time error correction (Ito, 2008; Popa & Ebner, 2019; Wolpert et al., 1998). In stuttering, cerebellar deficits may disrupt the formation of forward models, producing erroneous sensory predictions that trigger inappropriate error-correction responses, which in turn disrupt the coordination of speech. While atypical cerebellar volume and activity have been reported in PWS, their interpretations are not straightforward. Structural imaging studies suggested that anomalies in the cerebellum are associated with the etiology of stuttering (Chow et al., 2023; Chow & Chang, 2017; Connally et al., 2014). However, increased activity and functional reorganization in the cerebellum were also observed after speech fluency training, and cerebellar-frontal connectivity has shown to be negatively correlated to stuttering severity (De Nil et al., 2001; Sitek et al., 2016). These functional studies indicate that the cerebellum may play a compensatory role in stuttering, possibly compensating for basal ganglia dysfunction (Breska & Ivry, 2018; Grube et al., 2010; Kotz & Schwartze, 2010; Teki et al., 2011).

Most previous behavioral and neuroimaging studies employed case-control analyses at the group-level, which implicitly assume that stuttering is a homogeneous condition. To investigate inter-individual heterogeneity and potential neural subtypes in PWS, there is a need for alternative approaches that can identify brain atypicalities at the individual level. Moreover, because stuttering emerges in early childhood, a period of rapid brain development, characterizing individual structural and functional atypicalities in CWS requires the consideration of typical developmental trajectories.

Normative modeling has been used to account for age-related developmental effects in studies of individual-specific brain features (Bethlehem et al., 2022; Rutherford et al., 2023). Initially developed to characterize growth curves for morphological features such as height and weight in typically developing children, these methods are able to capture complex non-linear developmental trajectories (Borghi et al., 2006; De Onis, 2006). By modeling sex-specific brain development in typical children, normative modeling establishes a reference framework against which individual-specific neural atypicalities can be quantified. Individuals who exhibit similar patterns of neural atypicalities can then be grouped together into subtypes reflecting convergent neural profiles. Because these subtypes differ in their underlying neural bases, the associated behavioral phenotypes may also vary.

In this study, we performed normative modeling to identify individual-specific brain structure atypicalities using gray and white matter volumes defined from structural MRI scans obtained from a large mixed-longitudinal dataset of CWS and fluent controls aged between 3 and 12 (Chow et al., 2023). CWS were then clustered into subtypes based on similarity of individual atypicality maps. Neural subtypes will be further characterized on the basis of their behaviors and demographics. Previous group analyses of this dataset have been reported in Chow et al., 2023.

Based on prior literature and the framework proposed by Chang and Guenther (2020), we hypothesize that stuttering may arise from atypicalities in these three structural systems, (1) the BGTC loop, (2) the cerebellum, and (3) cortical speech-motor regions and associated white matter pathways that interface with the BGTC loop or cerebellum. Because cortical areas are generally more adaptable, we would expect that neuroanatomical differences in the basal ganglia and the cerebellum will be associated with higher stuttering severity than differences localized to cortical speech‐motor regions and their connections. Moreover, given the distinct functional roles of the basal ganglia and cerebellum in motor control, we anticipate that deficits in the basal ganglia will be associated with difficulties initiating and terminating speech sounds, leading to a higher frequency of dysrhythmic phonations, whereas cerebellar deficits result in an over‐reliance of feedback control and excessive corrective adjustments after speech initiation, leading to a higher frequency of repetitions.

## METHODS

### Participants

In this study, we performed secondary analyses on a large stuttering neuroimaging dataset used in our previous study, in which each pediatric participant was scanned up to four times at approximately one-year intervals (Chow et al., 2023). We included a total of 465 scans (240 CWS, 225 controls) after removing scans affected by head movements and other potential scanning issues. Stuttering severity was evaluated with the Stuttering Severity Instrument (SSI-4) (Riley, 2009). CWS were diagnosed with stuttering during their initial visit by considering their composite SSI scores (*>=*10), %SLD (*>*3 %), expressed concern of parent, and clinical confirmation. A child was considered recovered (i) if their composite SSI-4 was below 10 at their second visit or thereafter, (ii) percent occurrence of stuttering-like disfluencies in the speech sample was below 3, (iii) and after considering clinician and parent reports. In cases where their recovery status was not clear, a phone interview 1-2 years post final visit was used to track and evaluate recovery. Based on this method, 95 CWS were divided into 72 persistent CWS and 23 recovered CWS. For detailed demographics and behavioral characteristics of the participants, please review our previous publication (Chow et al., 2023). The Institutional Review Board at Michigan State University approved of all procedures in the study. Subjects’ consent was obtained in accordance with the Declaration of Helsinki. All parents signed a written informed consent, and all children either provided verbal assent (non-readers; typically, 3–5-year-olds) or signed a written assent form (readers; typically, 6 years and older).

### Image acquisition and preprocessing

Neuroanatomical images were acquired on a GE 3T Signa scanner (GE Healthcare) with an 8-channel head coil at Michigan State University. In each scanning session, a whole brain 3D inversion recovery fast spoiled gradient-recalled T1-weighted images with cerebrospinal fluid suppressed, was obtained with the following parameters: echo time = 3.8ms, repetition time = 8.6ms, inversion time = 831ms, inversion repetition time = 2332ms, flip angle = 8°, and receiver bandwidth = 620.8 kHz. These scans were collected as a part of a larger, ongoing longitudinal study, in which participants also underwent diffusion tensor imaging (DTI) and resting-state fMRI scanning.

Voxel-wise volumetric maps from the voxel-based morphometry (VBM) analysis from the previous study by Chow et al. were used as the input for this study. To summarize, a VBM analysis was conducted using the CAT12 toolbox. The study-specific templates based on a large publicly available data set and the demographic features (age and sex) of the participants was created using the CerebroMatic toolbox (Wilke, 2018). Volume of each voxel were obtained by multiplying (or modulating) tissue probability by the deformation field derived from the DARTEL normalization procedure (Ashburner, 2007). Individual, modulated images were resampled to 1.5 mm isotropic voxels and spatially smoothed with a Gaussian kernel with FWHM of 6 mm.

### Normative modelling and individualized centile maps

Gray and white matter volumetric maps from the VBM analysis were used to extract neural atypicalities for each CWS. A normative growth chart was generated using scans from control participants, to be used as a reference. Through a comparison to these reference growth charts, we quantified atypical brain volume for each stuttering scan. Each of these steps was done on a per-voxel basis. Refer to Figure 1 for a schematic of this workflow.

**Figure 1.**
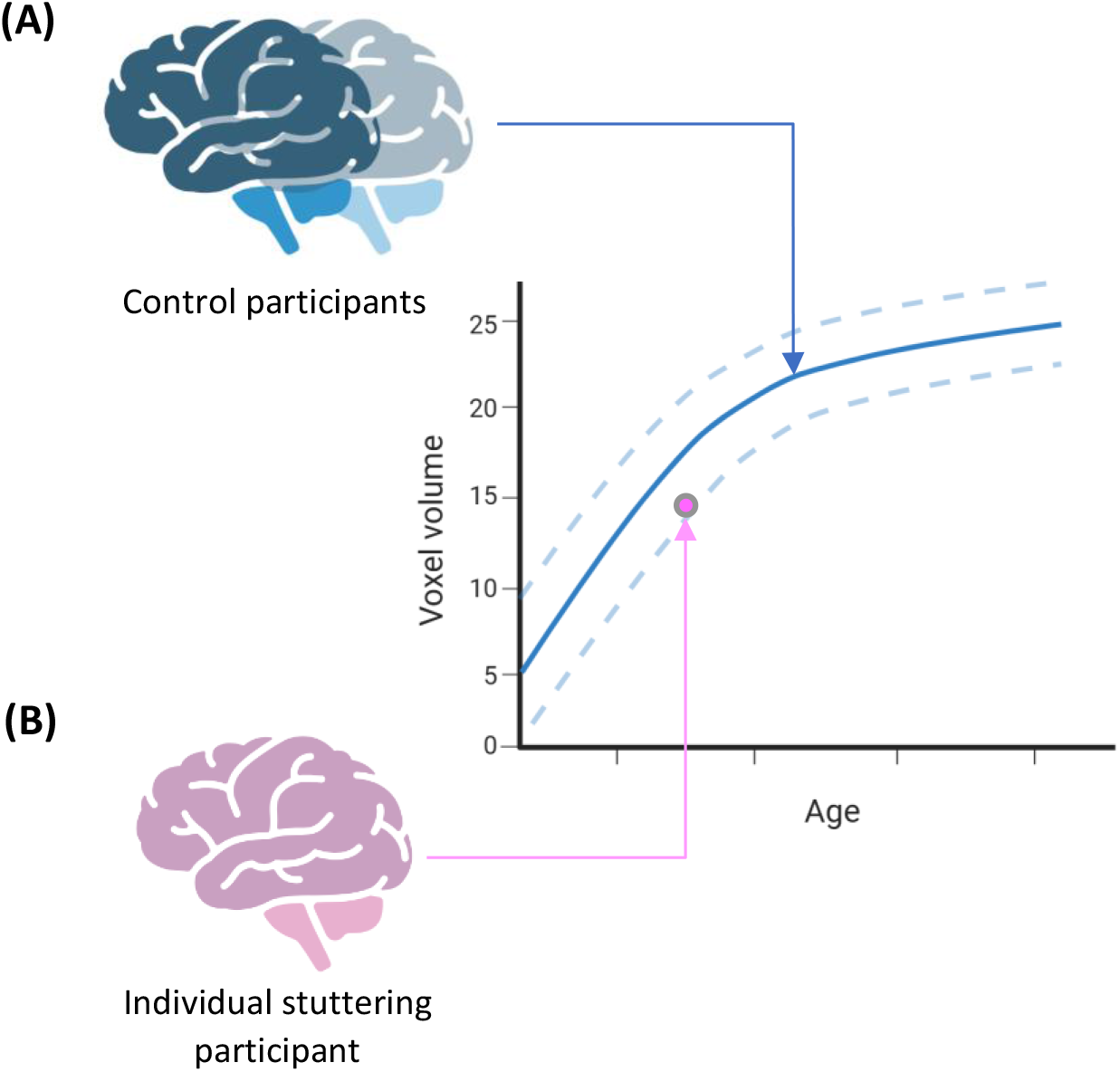
An illustration of normative modeling and the generation of individualized centile maps. (A) Voxel-wise volume maps from control participants were used to generate a normative growth curve for each voxel. (B) Voxel-wise volume of individual stuttering participants was compared with the normative growth curves at the participant’s age of scanning to calculate deviation in terms of centile for each voxel (individualized centile maps)

#### Model generation and specification

To conduct normative modeling, we used the generalized additive model for location, scale and shape (GAMLSS), a flexible, fast fitting univariate regression modeling analysis which allows exploration of large and complex datasets (Rigby et al., 2005). Unlike generalized linear model (GLM) and generalized additive model (GAM), GAMLSS allows the modeling of variance (σ), skewness (*ν*) and kurtosis (τ) independently in terms of explanatory variables. In contrast, GLM and GAM only model the variables’ means (μ), but not other statistical moments. In this study, modeling these additional variables allowed control over shape of the estimated developmental trajectories. The developmental trajectories of different morphometric measures (gray/white matter volume, cortical thickness, etc.) have previously been examined using multiple large neuroimaging datasets and GAMLSS (Bethlehem et al., 2022). In this previous study, the authors experimentally evaluated the optimal model specification and the underlying distribution based on Bayesian information criterion (BIC). In this current study, we implemented their findings for variable distribution and model specification for gray matter volume (GMV), subcortical gray matter volume (sGMV) and white matter volume (WMV). The model specifications for different tissue types are listed below.

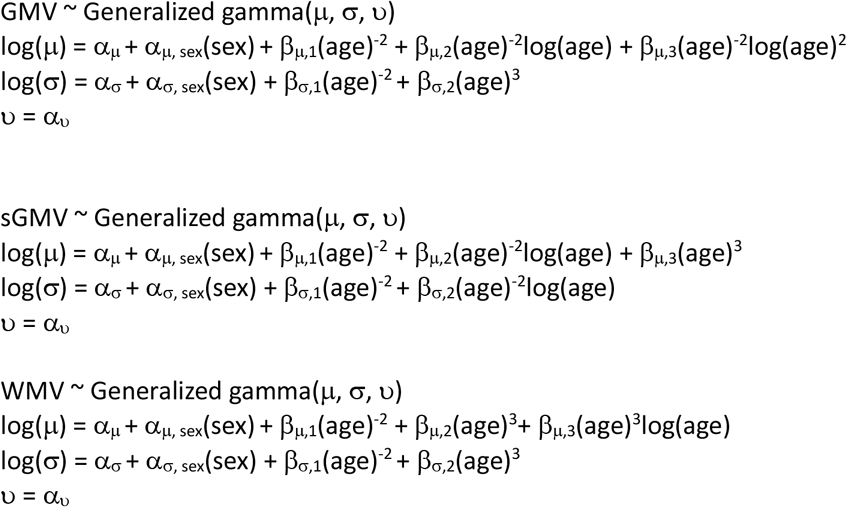

For each component of the generalized gamma distribution, α terms correspond to fixed effects of the intercept and sex; β terms correspond to fixed effect of the age modeled as fractional polynomial functions with the number of terms reflecting the order of the fractional polynomials. The link-functions for each component of the generalized gamma distribution have been explicitly mentioned.

To account for varying brain sizes, proportion volume maps were used for normative modelling. Specification and fitting of the GAMLSS model were implemented using the *gamlss* package in R.

#### Validation and reliability

In order to ensure that the reference norms generated from our control subjects were representative of typical populations, we checked whether we observed several key brain developmental trajectories previously found in typically developing children using the same model. Specifically, we examined the overall developmental trajectories of cortical gray matter, subcortical gray matter and white matter. In accordance with a study implementing the same technique in a large dataset, we would expect both sGMV and WMV to continue to increase through childhood while GMV would peak during early childhood.

To further study more fine grain cortical developmental trajectories, we found the age for peak GMV in each region defined by the AAL (automated anatomical labelling) atlas. The age at which the volume in a given ROI was maximum, was calculated by taking the median peak age across voxels within each ROI.

Model reliability and stability were evaluated through a bootstrap analysis. We ran 1,000 bootstrap iterations with stratified (5-fold) sampling with replacement to evaluate confidence intervals from the mean and standard deviation of the GAMLSS model. We observed a narrow confidence interval for both these terms. The bootstrap replicates were stratified by sex. To assess model goodness of fit, we examined detrended transformed Owen’s plots and Q-Q plots. Both these metrics showed that residuals were normally distributed.

#### Individualized centile maps

The goal of the normative modeling was to remove the typical developmental trajectories and highlight atypicality of gray and white matter volume in individual CWS. We used the models generated using control scans as a reference norm to calculate deviation in terms of centile in our age range and transformed voxel-wise the volumetric map of each CWS to a voxel-wise centile map.

### Clustering

#### Manifold, non-linear dimensionality reduction

After the generation of individualized centile maps, we performed clustering to identify CWS with similar patterns of neural atypicalities. We first used Isomap, a manifolding method to reduce the dimensions. Isomap performs non-linear dimension reduction by estimating the intrinsic geometry of the data.

In order to determine the appropriate number of dimensions the data should be reduced to, we needed to find a balance between low dimensionality such that the clustering algorithm would be effective and high dimensionality to retain as much information as possible. In order to take these two into consideration, we (1) calculated eigenvalues from the Isomap distance matrix (to identify the elbow point) and (2) determined the number of unclassified based on clustering of a default OPTICS algorithm. After balancing these two factors, we settled on reducing the dimensions to 5. Due to the high dimensionality of the initial data, we used Manhattan distance metric for this step. The k-nearest neighbors parameter was set to 45. It should be noted that the selection of k-nearest neighbors had to account for up to 4 scans from the same participant, hence it is higher than typical values.

#### Clustering algorithm

OPTICS (Ordering Points to Identify the Clustering Structure) was used for clustering. Similar to a closely related method, DBSCAN (Density-Based Spatial Clustering of Applications with Noise), OPTICS relies on density of data points, hence the geometric nature of Isomap worked well unison. Density-based algorithms group data points based on their proximity to dense regions of points. These dense regions of points, referred to as core points are identified, subsequently, directly and indirectly reachable points are identified. OPTICS differs from DBSCAN by allowing a variable radius for the identification of core points and reachable points. It maintains an ordering for how reachable points are, thus, the points are each assigned a reachability and ordering. These are used to define cluster boundaries.

From the OPTICS algorithm, we were able to extract reachability distances and ordering for each of the data points. Using the reachability and ordering for each point, we wrote a custom two-step script to identify clusters. In the first step, we identified when the slope for reachability sharply increases, indicating the start of the unclassified region. This point was also the boundary of the last cluster. In the second step, we followed a procedure similar to the ‘xi’ clustering of OPTICS, wherein we looked for valleys in the reachability, indicating regions of densely connected points. The parameters selected for OPTICS were min_samples=6 and the maximum steepness parameter used in clustering was 0.9. The *Scikit-learn* implementation of manifolding and OPTICS was used to implement these algorithms.

#### Representative neural map for each cluster maps

Once the group membership of each scan was established through OPTICS clustering, we generated a representative map for each cluster using raw volumetric data (without normative modelling). In order to do this, we trained a mixed effects model for each voxel, with volume as the dependent variable and sex, cluster membership, age, quadratic age as independent variables. The intention here was to utilize the main effect of cluster membership to identify voxels that are atypical in a given cluster when compared to control participants. A mixed-effects model was needed to account for multiple measurements from a single individual. We also included IQ, socio-economic status, total intracranial volume and SSI as covariates.

Using the coefficients from the mixed effects model, we generated a neural map with the t-value for each cluster; thus, for each voxel, we calculate a weighting associated with the given cluster, generating a representative map for the cluster. Voxel-wise height threshold of p < 0.005, and cluster-size thresholds of k > 457 voxels in GM and k > 318 in WM were used to control for false positives. This set of thresholds is equivalent to correct p< 0.05. It is important to mention here that we used proportion maps for the normative centile mapping and subsequent clustering, but after identifying the group membership, we switched to raw volume maps when generating the representative map for each cluster. For the cluster-specific maps, we used raw maps in the linear mixed effects model because raw VBM data is a closer reflection of brain volume and the total volume is controlled for using total intracranial volume.

### Statistical testing of behaviors and demographics

Behavioral and demographic characteristics of sub-group clusters were tested by making comparisons between each pair of clusters. SSI, SSI sub-scores (frequency, duration, secondary characteristics) and disfluencies (repetitions, dysrhythmic phonations, repeated utterances, typical disfluencies) were tested using logistic regression models with age, sex, socioeconomic status and IQ as covariates. SSI and the frequency sub-score were each tested using a separate model with these covariates. A third model was specified including duration, secondary characteristics, and disfluencies and the covariates. Additionally, recovery was added as a binary variable. SSI and frequency could not be included in this larger model because they showed a high correlation with other variables, thus not allowing proper model fitting.

Differences in the ratios of sex (females, males) and recovery (persistent, recovered) were tested using a Fisher Exact test. Statistical functions were implemented using the python *Scipy* module and the logistic regression was implemented using the *Statsmodels* module in python.

## RESULTS

### Normative growth charts

To measure individual-specific atypicality in brain volume in CWS, we generated reference normative growth charts using scans from control participants. To validate these growth charts, we qualitatively compared them to growth charts from other studies that investigated typical development in the childhood age range (Bethlehem et al., 2022; Giedd et al., 1999, 2015; Gogtay et al., 2004; Lenroot & Giedd, 2006; Raznahan et al., 2014; Tiemeier et al., 2010). When examining each tissue type across the whole brain, we found that gray matter volume increases during early childhood, reaching a peak at around age 5, before gradually decreasing (Figure 2A). In contrast, white matter and subcortical gray matter volume continue to increase into adolescence. Peak ages reported in this study aligned with Bethlehem et al., 2022, which implemented the same normative modeling technique to study brain development.

**Figure 2.**
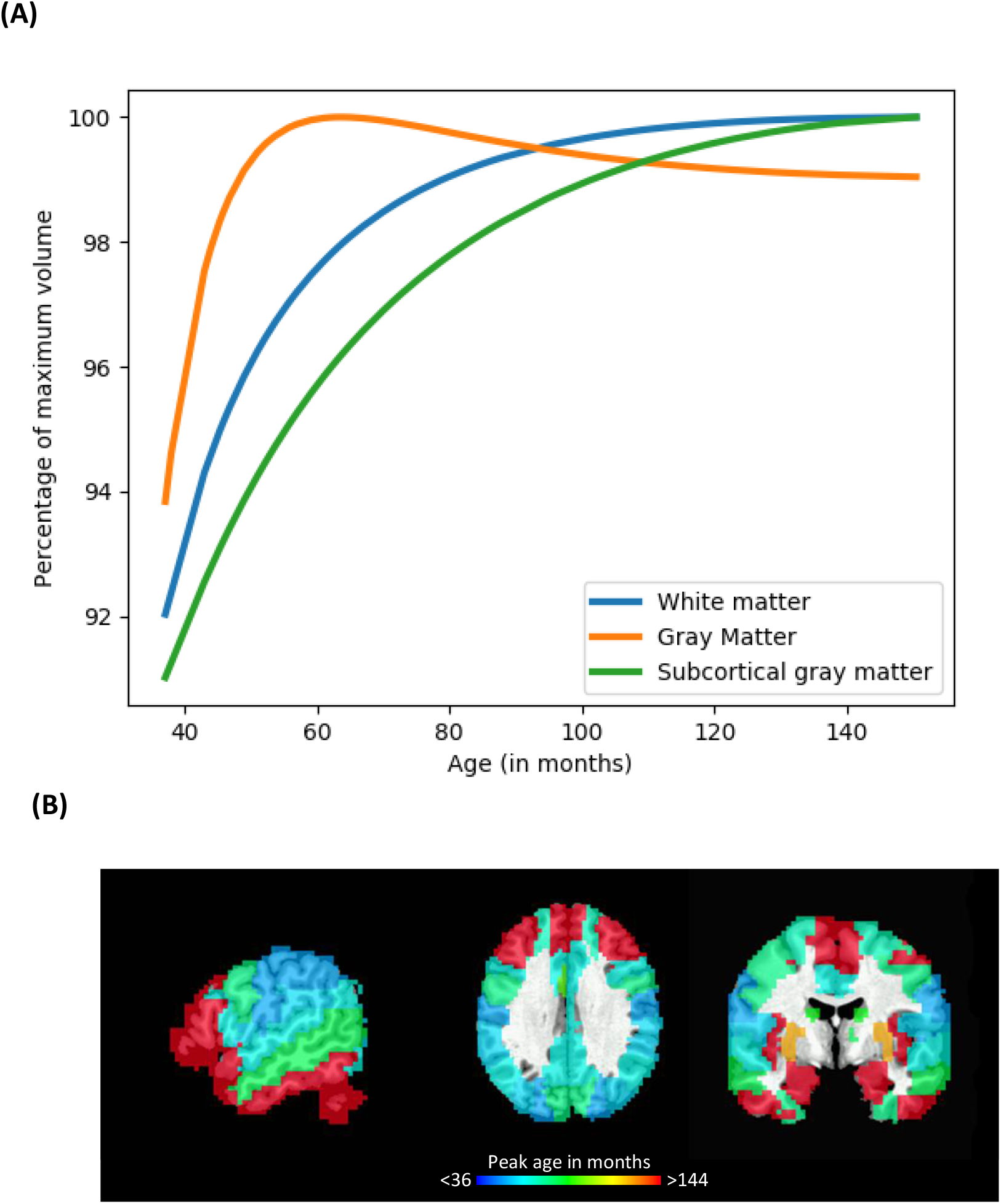
Brain volume development in control participants. (A) Averaged volume across voxels by tissue type, normalized by each tissue type’s maximum value across our sample’s age range. On average, cortical gray matter peaks at around 5 years of age and then decreases (orange line), while white matter (blue line) and subcortical gray matter (green line) continue to increase into adolescence. (B) The median age at which gray matter peaks across voxels in each cortical and subcortical areas defined by the AAL (automated anatomical labelling) atlas is overlaid on three selected slices of an anatomical image in MNI space ([x, y, z]: [-54, -6, 30]). The results are consistent with a previous study showing the motor cortex volume peaks later than the somatosensory cortex, and the anterior frontal regions peak relatively late in childhood (Bethlehem et al., 2022).

In a more fine-grained analysis of cortical gray matter volume developmental trajectories, we observed that the volumes in sensory regions generally peaked earlier in age than those in the frontal regions (Figure 2B). For example, the somatosensory cortex peaks earlier than the motor cortex. In the temporal lobe, the superior temporal gyrus (STG) peaks later than the middle temporal gyrus (MTG), which in turn peaks later than the inferior temporal gyrus (ITG). Striatal and thalamic volume curves flattened around 7-9 years of age. While there is some variability within cerebellar subregions, volume is generally found to increase into late childhood. Although developmental trajectories have varied across studies, the patterns observed in the present study were largely consistent with previous reports (Bethlehem et al., 2022; Giedd et al., 2015; Gogtay et al., 2004; Raznahan et al., 2014; Tiemeier et al., 2010).

### Individualized centile maps

Each scan from a CWS was compared to the reference growth charts to calculate deviation in terms of centile values. We observed that individualized maps from the same participant at different ages, closely resemble each other (Figure 3A, B), whereas maps between participants show greater variation (Figure 3B-D). We also observe similarities and differences across individuals. For example, both SM101 and SM133 show reduced cerebellar volume (Figure 3B, D), while centile values in the putamen are increased in SM101 but decreased in SM133.

**Figure 3.**
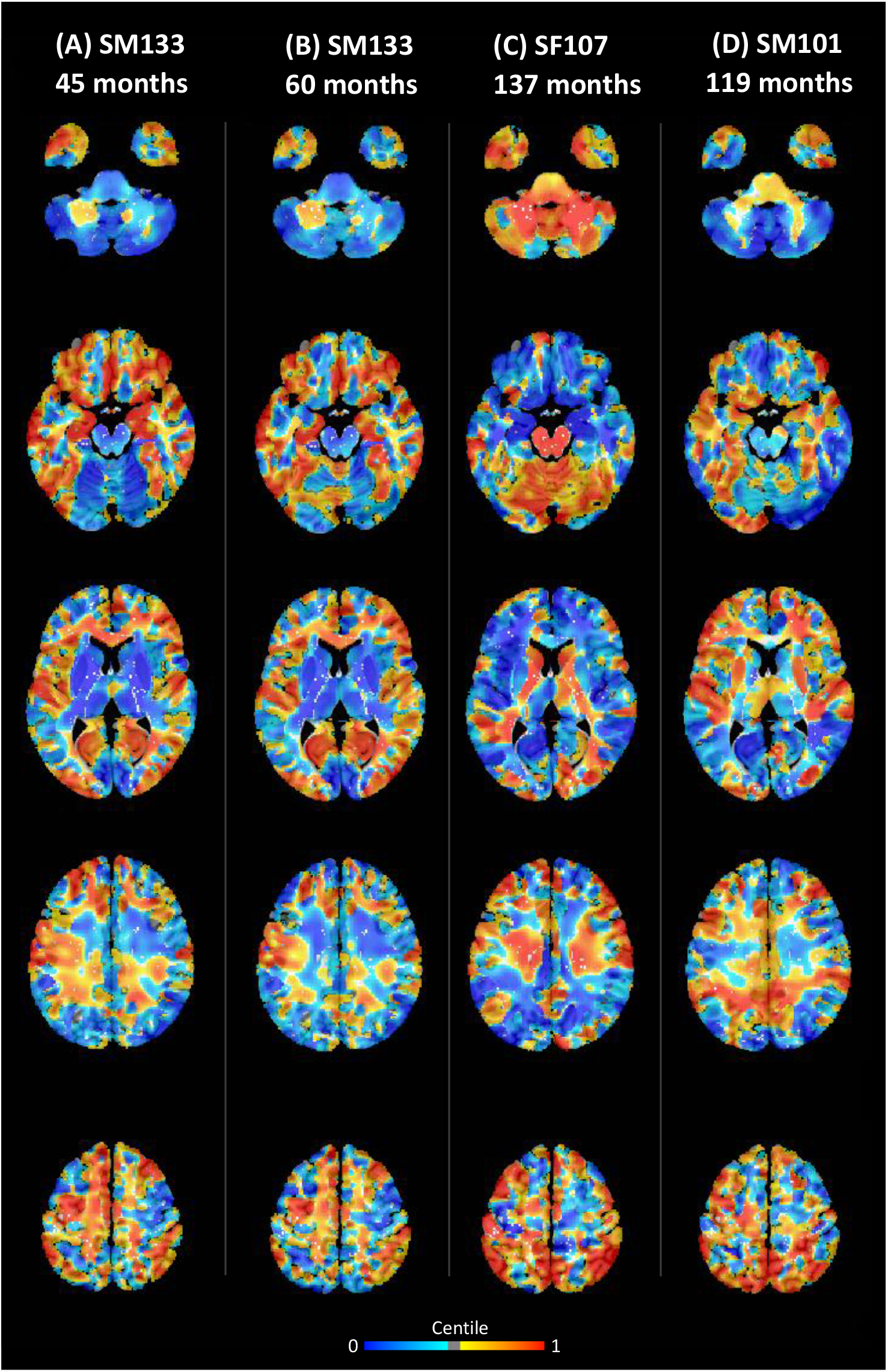
Examples of individualized centile maps of CWS. The voxel-wise deviations from the normative map in centile of three participants (A/B, C & D) are overlaid on selected axial slices of an anatomical image in MNI space. Warm colors indicate volume values greater than 0.5 centile and cool colors indicate values less than 0.5 centile. Centile maps (A) and (B) are from the same participant, at different time points. In general, centile maps from the same participants at different time points are relatively similar. Noticeable inter-individual differences are observed. For example, cerebellar volume of SF107 is relatively large compared to that of SM133 and SM101 (first row of images).

### Clustering results

We performed a data reduction step followed by a data-driven clustering algorithm to identify groups of individual CWS with similar individualized centile maps. These groupings or clusters are referred to as neural subtypes of stuttering. Overall, 192 scans (80%; or 86.32% of CWS) were grouped into four clusters, and 48 scans (20%; or 13.68% of CWS) were considered “unclassified”. Scans from a single CWS were grouped into a single cluster at most, in other words, there are no individuals grouped into multiple clusters. For the four identified clusters, each was characterized by distinct patterns of brain atypicality (Figure 4). Regions of atypicality for each cluster are described below.

**Figure 4.**
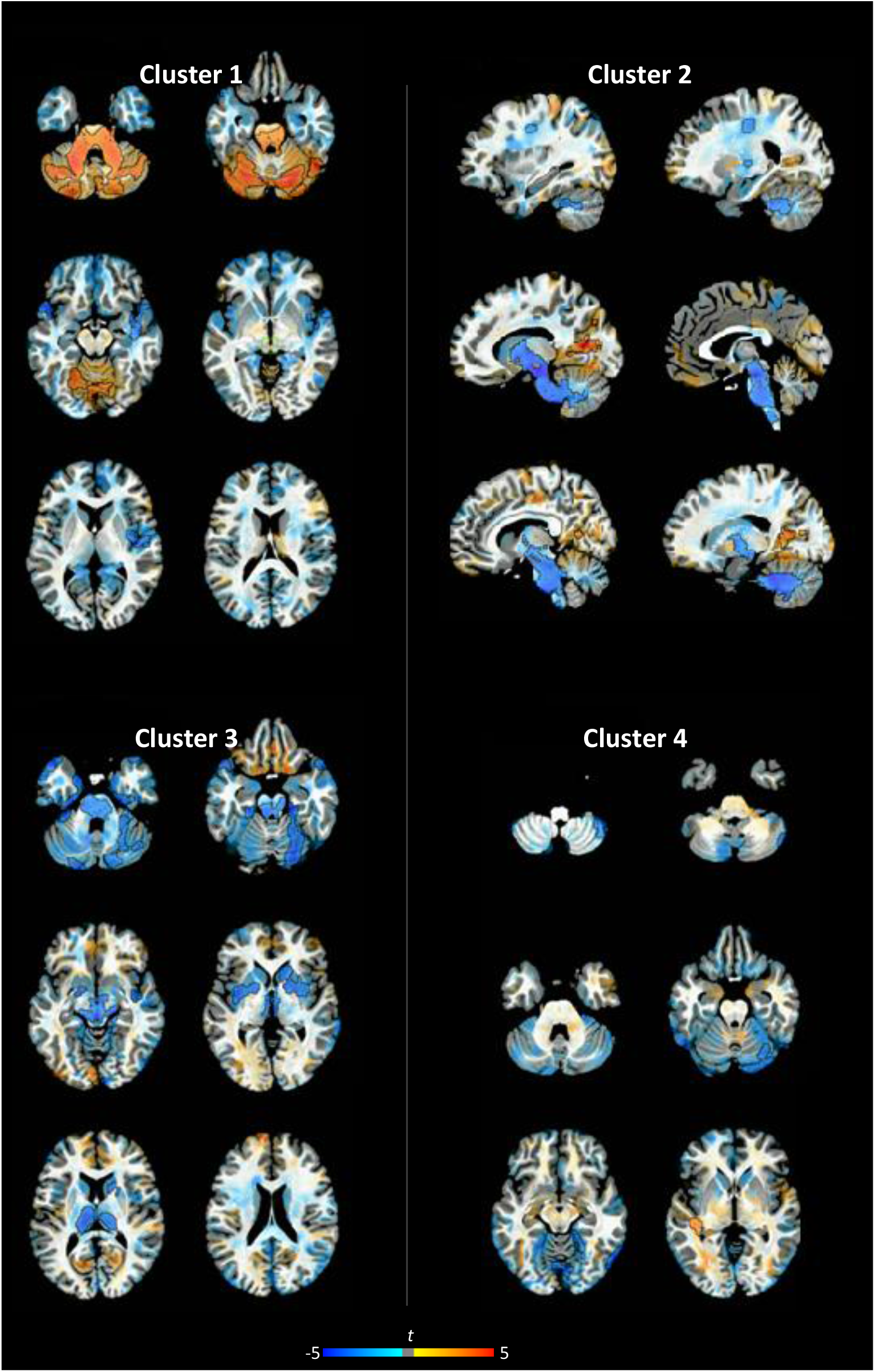
Differences in volume between each subtype cluster and controls using linear mixed-effects models. Group differences in *t*-statistics are overlaid on selected slices of an anatomical image in MNI space. Orange and blue indicate increased and decreased volume at uncorrected *p*<0.005, respectively. Clusters exhibiting a significant difference at corrected *p*<0.05 are outlined by black lines.

*Cluster 1* showed a marked increase in bilateral cerebellar volume, including both gray and white matter areas. Gray matter increases are predominantly in the lateral cerebellum. The increases in cerebellar volume were accompanied by decreased volume in insula and the superior temporal gyrus in the right hemisphere as well as the left superior temporal gyrus.

*Cluster 2*, the most notable feature of this cluster was a decrease in volume in a large white matter region spanning from the cerebellar peduncle, the brain stem, and the internal capsule, along with decreases in white matter tracts beneath the left sensorimotor regions. In addition to decreases, an increase in volume was observed in the bilateral visual cortex.

*Cluster 3* is characterized by reductions in gray matter volume in the BGTC loop, including the bilateral thalamus, caudate and putamen, as well as the anterior cerebellum, inferior temporal gyrus, and right superior parietal cortex.

*Cluster 4* showed volume reductions in the right lateral cerebellum, accompanied by an increase in the left temporo-parietal junction/planum temporale.

### Behavioral results

In addition to patterns of neural atypicality, each cluster was further characterized based on available behavioral measures and demographic features (Table 1 and Figure 5). Logistic regressions were used to find significant differences in behavioral measures between each pair of clusters. Cluster 3 was the most severe with an SSI of 20.93 and high values for the SSI sub-scores, frequency and duration. Further breaking down SLDs into different types, Cluster 3 showed a higher number of repetitions. The recovery rate was also low, with only 1 out of 18 children (5.6%) recovering from stuttering. Cluster 2 exhibited the mildest stuttering severity, with an SSI score of 15.34, consistent with relatively low SSI sub-scores and SLD rate, and the highest rate of recovery. The SSI scores of Clusters 1 and 4 were around the average SSI scores of all the CWS participated, with 17.99 and 18.53 respectively. Although similar in SSI scores, they differed based on behavioral measures. Cluster 1 showed higher SLD rate, whereas Cluster 4 showed a relatively low SLD rate but the highest scores for physical concomitants and number of dysrhythmic phonations.

**Table 1.** Characteristics of subtype clusters.

|  | Cluster 1 | Cluster 2 | Cluster 3 | Cluster 4 |
| --- | --- | --- | --- | --- |
| Number of individuals | 20 | 28 | 18 | 16 |
| Number of scans | 49 | 62 | 44 | 37 |
| Age at scan (months) | 85.86 | 65.36 | 78.31 | 98.26 |
| Sex ratio (boys:girls) | 9:11 | 18:10 | 13:5 | 10:6 |
| Recovery rate | 5/20 | 10/28 | 1/18 | 4/16 |
| **SSI** | 17.99 | 15.34 | 20.93 | 18.52 |
| Frequency | 8.62 | 8.04 | 9.81 | 7.69 |
| Duration | 5.99 | 5.59 | 7.13 | 5.72 |
| Secondary | 3.01 | 1.7 | 3.32 | 5 |
| **SLD counts per syllable** |  |  |  |  |
| Repetitions | 2.92 | 2.7 | 4.58 | 2.15 |
| Dysrhythmic phonations | 1.85 | 1.25 | 1.31 | 2.11 |
| Typical dysfluencies | 4.4 | 4.26 | 4.66 | 3.96 |
| Repeated Utterances | 1.16 | 1.22 | 1.25 | 1.19 |

**Figure 5.**
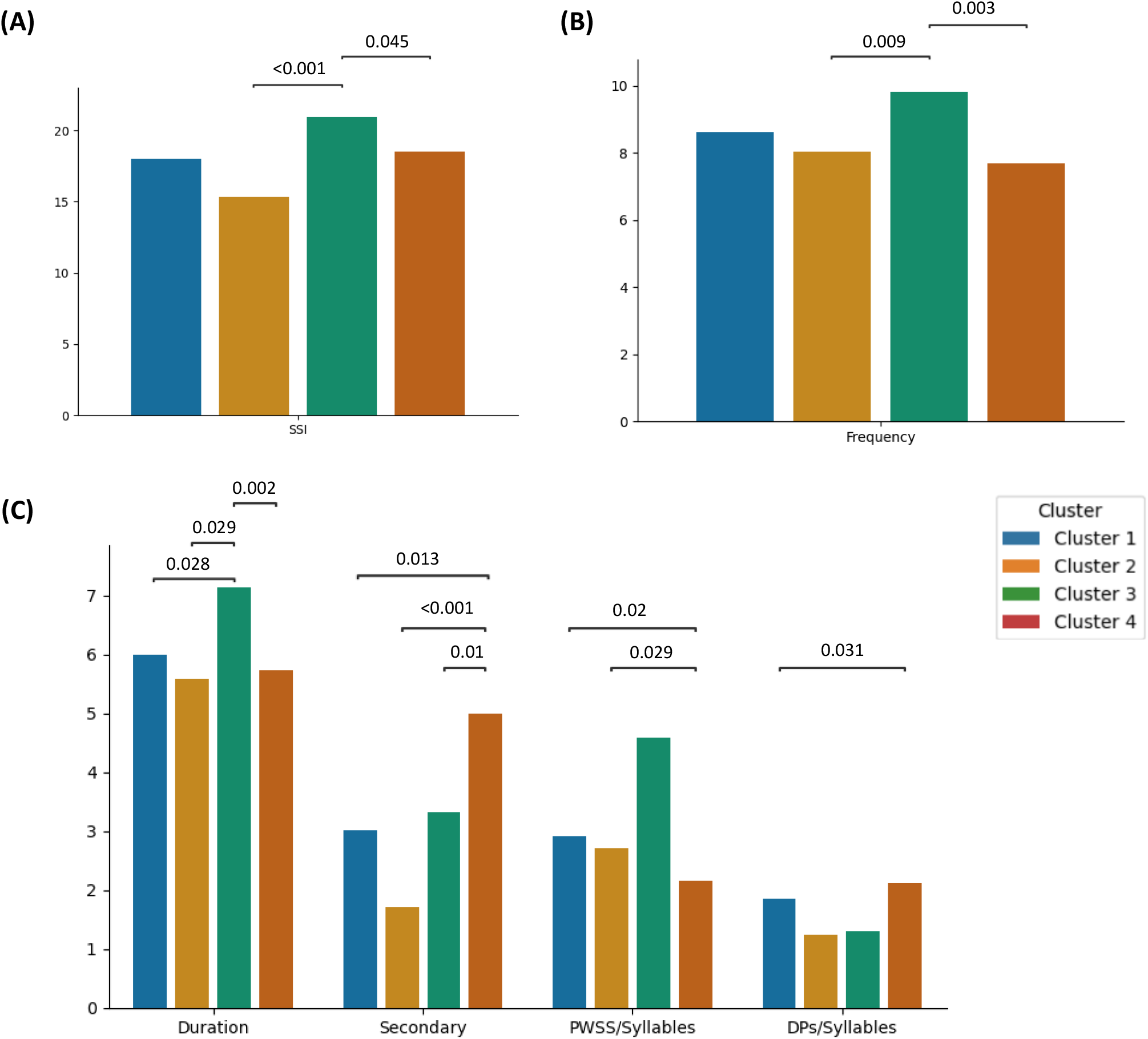
Behavioral metrics, stratified by subtype clusters. Three sets of logistic regression models were used to examine the contribution of behavioral metrics in separating a pair of subtype clusters: (A) SSI, (B) SSI frequency sub-score. (C) SSI duration and secondary sub-scores, part-word or single-syllable repetitions per syllable and dysrhythmic phonations (blocks and prolongations) per syllable. Typical dysfluencies and repeated utterances were also included in the model; however, no significant differences were identified for these variables. Statistically significant pairwise differences are annotated with p-values.

Cluster 1 had a higher proportion of girls compared to the other clusters (Cluster 1 > Cluster 2, p value = 0.003; Cluster 1 > Cluster 4, p value = 0.022), but the recovery rate was not significantly higher. The recovery rate of Cluster 2 was significantly lower than that of Cluster 3 (p value = 0.002).

## DISCUSSION

Inter-individual heterogeneity in stuttering behavior has been well documented, reflecting a complex neural basis shaped by the interplay between genetic and environmental factors (Smith & Weber, 2017). Isolating individuals with similar underlying neural mechanisms will be important for advancing our understanding of the disorder’s etiology. However, separating CWS who differ on the basis of underlying neural mechanisms is challenging; in this study we use morphological evidence in the form of brain volume to address this problem. Gray and white matter volumetric data were used to characterize individual-specific differences, or neural atypicalities. We then clustered individuals based on neural atypicalities to identify subgroups of CWS, termed subtypes.

To examine individual-specific differences, we compared voxel-wise brain volume of each CWS to a reference norm, thus, removing typical developmental trends to highlight individual-specific atypicalities. A prerequisite for this method is to generate a reliable norm that captures typical development. We qualitatively compared the developmental trends derived from controls with the results from a prior study which examined brain development in a large cohort of control participants using the same method (Bethlehem et al., 2022). Consistent with this prior study, our norm showed that cortical gray matter volume peaks at 5-6 years of age and both white matter and subcortical gray matter volume continue to increase in our participants’ age range (Figure. 2B). Additionally, in our analysis of regional gray matter development, we observed that cortical gray matter volume in posterior regions peaked at earlier ages than in anterior regions. This posterior to anterior pattern of brain development has been reported previously and is considered an index of cortical maturation (Gerván et al., 2017; Gogtay et al., 2004). The frontal lobe, for instance, is known to reach its peak volume later in childhood, consistent with that fact that executive functions mature at a later stage (Arain et al., 2013; Best & Miller, 2010). Striatal and thalamic volume curves flatten around 7-9 years of age. While there is some variability within cerebellar subregions, volume generally increases into late childhood. Although previous studies have reported different estimates of regional gray matter volume development depending on the modeling approaches (Giedd et al., 1999, 2015; Lenroot & Giedd, 2006; Raznahan et al., 2014), our results closely resemble the findings reported in a prior study using the same model (Bethlehem et al., 2022), suggesting that our normative model was implemented appropriately.

The cluster analysis classified 86% of the CWS into four neural subtypes with distinct neuroanatomical differences compared to fluent controls. In general, the regions identified in the subtypes appear to concentrate in the cerebellum and the BGTC loop, including associated white matter pathways. Our analysis showed that about 20% of CWS showed clear morphological differences in the basal ganglia proper (Cluster 3). Moreover, we did not observe clear evidence of subtypes based on different components of the BGTC loop as hypothesized in Chang and Gunther (2020). Instead, CWS seem to exhibit a wide range of differences in the cerebellum, and its associated cortical and white matter areas (Clusters 1, 2 and 4). It is possible that cortical and white matter alterations in these subtypes influence BGTC loop function, while the accompanying cerebellar alterations may represent compensatory neurodevelopmental adaptations.

The presence of cerebellar volume atypicalities across all four subtypes is perhaps the most critical finding in this study. Among the subtypes, cerebellar volume atypicalities varied in both anatomical location and direction of deviation (increases or decreases in volume relative to controls). For example, in Cluster 1, we observe an increase in volume in the cerebellar peduncles, whereas in Cluster 2, we observe decreases in volume in the same region. Such contrasting patterns of brain development would likely get obscured in a group-level analysis, wherein averaging across heterogeneous profiles may obscure meaningful differences. In addition to the direction of deviation, the location for cerebellar atypicalities varies per subtype, underscoring variability in how the cerebellum may be implicated in stuttering.

The cerebellum supports speech production by implementing forward models that generate predictive estimates of the sensory consequences of planned articulatory movements (Guenther, 2019; Ito, 2008; Wolpert et al., 1998). These predictive computations allow the cerebellum to refine and coordinate motor commands through error-based updating, supporting the precise timing and fluent execution of speech (Ackermann, 2008; Sokolov et al., 2017). Prior work has shown that stuttering persistence is linked to functional and structural differences in the cerebellum, and PWS exhibit heightened cerebellar activation alongside reduced structural connectivity between cerebellar and motor cortical regions (Connally et al., 2014; De Nil et al., 2003; Lu et al., 2010; Neef & Chang, 2024; Watkins et al., 2008). On the other hand, cerebellar activation has been linked to fluent speech production and increased cerebellar peduncle growth has been associated with reduced stuttering severity (Chow et al., 2023; Fox et al., 2000). These increases have been interpreted as reflecting cerebellar compensation, which attenuate stuttering severity but may be insufficient to support complete recovery (Alm, 2004; Chow et al., 2023; Kotz et al., 2009; Petacchi et al., 2005). These findings paint a complex picture of the cerebellum’s structural and functional involvement in stuttering, spanning its role in development, persistence, recovery, and the support of speech fluency. In the following paragraphs, we will discuss each subtype, highlighting atypical regions identified and noting potential interactions with the cerebellum.

In Cluster 3, we observe prominent decreases in volume in the components of BGTC loop, including the putamen, caudate, and thalamus as well as the cerebellum, in CWS compared to controls. This cluster is consistent with contemporary neuroanatomical models of stuttering (Chang et al., 2019; Neef & Chang, 2024). In speech-motor control, the BGTC loop plays important roles in sequencing and initiation of movement (Bohland et al., 2010; Civier et al., 2013). The basal ganglia are considered a component of the feedforward control system, receiving inputs from the speech sound map, initiation map, and articulator map.

This subtype presented the most severe phenotype in terms of stuttering severity, as measured by SSI-4, and lowest rate of recovery. It also showed a higher frequency of SLDs driven by repetitions, and longer duration of stuttered syllables. Gray matter abnormalities in both the BGTC loop and the cerebellum may be a marker for higher severity and higher likelihood of persistence. Taken in conjunction with the neuroanatomical findings, we speculate aberrant timing signals from the basal ganglia may lead to incorrect syllable initiation and sequencing, and overcorrection in cerebellum in terms of both timing cues and sensory feedback may lead to disfluencies in this subtype.

The low rate of recovery compared to other subtypes suggests that this subtype may be more prevalent in adults compared to children. Due to the higher prevalence in adults and higher stuttering severity, this subtype may represent a well-studied sample of individuals, resulting in it being considered a prototypical neural model of stuttering.

The most prominent feature of Cluster 2 was a large region of decreased white matter volume spanning the cerebellum, brain stem, and internal capsule, and reduced white matter volume in tracts underlying left sensorimotor regions. These white matter pathways are likely corticocerebellar and motor descending pathways, which are critical pathways for motor planning and control. However, these pathways may represent direct cerebellar-basal ganglia projections; while traditional models suggest that interactions between the cerebellum and basal ganglia are mediated by the cortex and thalamus, recent studies have found direct tracts connecting the cerebellum and basal ganglia (Bostan et al., 2010; Hoshi et al., 2005; Milardi et al., 2016; Quartarone et al., 2020). These direct cerebellar-basal ganglia tracts, in turn, give the cortex the ability to fine-tune decision making and motor action selection (Quartarone et al., 2020). Additional tractography studies will be needed to further study the nature of these white matter atypicalities.

This subtype showed the mildest stuttering severity scores, and no other behavioral characteristic stood out. This subtype showed the highest rates of recovery. White matter-based neuroplasticity allows for the acquisition of new motor skills (Fields, 2015; Sampaio-Baptista et al., 2013). This compensatory rewiring of white matter connections may account for the relatively mild phenotype and higher rates of recovery in this subtype.

Cluster 1 and 4 did not show basal ganglia abnormalities, however, both these subtypes exhibited prominent cerebellar atypicalities. In Cluster 1, increased cerebellar volume was accompanied by decreases in volume in the right insula. The insula receives input from somatosensory and auditory cortices and cross modal processing of sensory information might be a prerequisite for fluent speech production (Calvert, 2001). The insula has been associated with an auditory-phonological system that provides sensory targets for the vocal tract (Hickok et al., 2011). The insula also has reciprocal connections with the orbitofrontal cortex and the anterior cingulate cortex (ACC), and these are structures involved with the control of emotional and motivational states (Ackermann & Riecker, 2004; Augustine, 1996). At the junction of the affective and motor control networks, the insula is suited to add affective-prosodic information to verbal utterances during speech production. Through efferent connections to the premotor cortex, the insula has critical sensorimotor contributions which may then have a downstream role in cerebellar motor control. Moreover, both the insula and cerebellum are engaged in facilitating speech on a moment-to-moment basis, moments of disfluency may arise from imbalances in on-line processing of speech.

Cluster 1 exhibited a mild/moderate severity phenotype and interestingly included a greater proportion of females relative to the other subtypes. A previous fMRI study in AWS found sex-specific differences in the right insula during speech planning, with decreased activation in the stuttering group driven by females (Chang et al., 2009). This suggests that structural and functional abnormalities in the insula may be present in a subset of PWS.

In Cluster 4, decreases in volume were observed in the right cerebellum and increases in volume were observed in the left temporo-parietal junction (TPJ), including the planum temporale. Being at the junction of auditory, somatosensory, visual cortical regions, the TPJ is known to play an important role in sensorimotor integration. This region contributes to generating sensory representations of speech sounds. It has previously been speculated that these representations are accurately coded in PWS, however, the mapping between the sensory and motor representations may be imprecise (Hickok et al., 2011). As a result of the poor mapping of the internal model to the sensory system in the TPJ, incorrect predictions may be generated in the cerebellum and error correction in turn will be inaccurate. We speculate that inaccuracies in the sensory-motor mapping contribute to stuttering in this cluster.

Similar to the previously subtype, this subtype exhibited a mild/moderate severity phenotype. However, this subtype showed a high number of dysrhythmic phonations and physical concomitants. Compared to other subtypes, severity in this subtype does not appear to be driven by repetitions.

To conclude, we provide an overview of the neural mechanisms associated with each of the four subtypes. BGTC impairments in Cluster 3 suggest deficits in the feedforward control of speech and aberrant timing may lead to incorrect speech syllable initiation and sequencing. Cluster 1 and 4, both show cerebellar alterations alongside abnormalities in regions involved in auditory-motor mapping. We speculate that there are deficits in sensorimotor integration and error correction in these subtypes. Finally in Cluster 2, we find deficits in descending motor white matter pathways suggesting irregularities in motor execution.

Several limitations of this study should be noted. Although this represents the largest neuroimaging dataset of stuttering to date, the sample size remains modest for this analytical approach. Statistical power is further reduced when the cohort is divided into subtypes, limiting sensitivity for between-subtype comparisons. Another key limitation is the recovery rate in the sample is relatively low (24%), when compared to commonly reported estimates (80%). This discrepancy is likely due to recruitment of older children for feasibility in a neuroimaging study. While the cohort included younger children, many participants were recruited after the age five, at which point the likelihood of stuttering persistence increases. As a result, the cohort included children more likely to persist stuttering, limiting the generalizability of recovery rates. Additionally, behavioral characteristics compared in this method were limited to SSI subscores and disfluency types, thus, the scope of behavioral characterization was relatively narrow. Including other measures such as performance in rhythm, sensorimotor adaptation, and motor learning tasks would improve the ability to link neural differences to broader functional profiles.

Withstanding these limitations, examining individual-specific differences and subtyping based on neural atypicalities provides valuable insight and represents a promising future direction. Herein, we highlight the importance of investigating neural heterogeneity in developmental stuttering. The most salient findings from the subtyping were the identification of a basal ganglia-thalamo-cerebellar subtype with higher severity and lower rates of recovery, and a white matter subtype with mild severity and a higher likelihood of recovery. Findings from this study suggest that cerebellar contributions to stuttering warrants further examination. Further studies are also needed to investigate the functional consequences of structural atypicalities identified in this study. The normative modelling implemented in this study, could be expanded to incorporate other MRI modalities, such as resting state functional connectivity, or biological measures, including physiological recordings or genetic information, to better characterize the subtypes.

An important direction for future work is to incorporate scans from additional sites into the normative modeling and subtyping analysis. The primary challenge would be to account for inter-site variability to harmonize the different datasets. Nonetheless, through the inclusion of a greater number of scans from different sources, we could verify the robustness and stability of the subtypes. The neural subtype framework developed here also holds potential clinical application. Children scanned at any site could be classified into subtypes, enabling interventions tailored to neural profiles. This approach could support more personalized treatment by linking subtypes to behavioral characteristics and treatment responsiveness. However, substantial work is still needed to validate the subtypes across larger cohorts and to assess clinical outcomes beyond recovery, including responsiveness to different treatments for each subtype

## ACKOWLEDGEMENTS

The work was supported by the National Institute on Deafness and Other Communication Disorders (NIDCD), National Institutes of Health under award numbers, R01DC020157 (PI: Chow) and R01DC011277 (PI: Chang). The content is solely the responsibility of the authors and does not necessarily represent the official views of the NIDCD or the National Institutes of Health.

